# A novel core promoter element induces bidirectional transcription in CpG island

**DOI:** 10.1101/202622

**Authors:** Amin Mahpour, Dominic Smiraglia, Benjamin S. Scruggs, Irwin H. Gelman, Toru Ouchi

**Affiliations:** Department of Cancer genetics and Genomics, Roswell Park Cancer Institute, Buffalo, NY, 14263, USA; Epigenetics and Stem Cell Biology Laboratory, National Institute of Environmental Health Sciences, Research Triangle Park, NC, 27709, USA

**Keywords:** Transcription, Bidirectional Transcription, Gene Regulation, CpG Islands, TATA-less Promoter, Core Promoter Element

## Abstract

How TATA-less promoters such as those within CpG islands (CGI) control gene expression is still a subject of active research. Here, we have identified the “CGCG element”, a ten-base pair motif with a consensus sequence of TCTCGCGAGA present in a group of promoter-associated CGIs of ribosomal protein and housekeeping genes. This element is evolutionarily conserved in vertebrates, found in DNase-accessible regions and employs RNA polymerase 2 to activate gene expression. Through extensive analysis of several endogenous promoters, we demonstrate that this element activates bidirectional transcription through divergent start sites. Methylation of this element abrogates the associated promoter activity. When coincident with a TATA-box directional transcription remains CGCG-dependent. Because the CGCG element is sufficient to drive transcription, we propose that its unmethylated form functions as a core promoter of TATA-less CGI-associated promoters.

## Introduction

Gene expression is one of the most critical, yet enigmatic, biological processes that defines cellular and organismal identity, and that mediates cellular response to internal and external stimuli ^1^. Importantly, dysregulation of this process is known to contribute to various human diseases such as cancer ^2^. With the discovery of RNA polymerases, the mechanisms of how transcription occurs have been extensively studied in many organisms ^3^. In contrast to the relatively simple prokaryotic transcriptional system, metazoan transcription is considerably more elaborate and involves complicated promoter structures, multiple functional DNA elements and a repertoire of specific general transcription factors. These factors and DNA elements are required to facilitate accurate transcriptional initiation, elongation, and termination ^4–6^.

The best-known DNA element that mediates the initiation of transcription of protein-coding genes is the TATA box with the consensus sequence TATAA ^7^. This element is usually located 25 to 34 base pairs upstream of transcription start sites (TSS). However, most human promoters, including those regulating housekeeping genes lack this DNA element ^8^, suggesting that TATA-less promoters are controlled by different yet poorly understood mechanisms. A few novel elements have been described that presumably function as core promoter elements in TATA-less promoters ^9–12^. Yet, most of these promoter elements (e.g. GC-box or Inr motif) require additional transcriptional activator binding sites in order to drive directional transcription.

Vertebrate genomes contain short G+C rich sequences that are typically less than 1 kb long traditionally termed CpG islands (CGIs) ^13, 14^. These regions are considered to be critical for transcriptional regulation of a large group of genes that include housekeeping genes ^15^. Most CGI-associated promoters lack a TATA box yet contain “GC-box” binding sites for the general transcription factor SP1 although GC box is not sufficient to induce transcription on its own ^15–18^. CGI-associated promoters typically induce bidirectional transcription that produces coding and non-coding transcripts ^19,20^. Thus, depending on the stability of the non-coding RNA, CGI-associated promoters can generate more stable long non-coding RNAs (lncRNA) or short-lived transcripts ^21^. To date, no specific independently-acting promoter element governing these CGI-associated bidirectional promoters has been described.

In this study, we analyzed DNase accessible CGIs in the K562 cell line and found an enriched motif with the consensus sequence of TCTCGCGAGA, which we termed the “CGCG element” due to the characteristic central bases. This element confers transcriptional activity independent of other transcriptional activator sequences. Promoter sequences related to the CGCG element have been reported previously for several individual genes, but their functional significance was never explored ^22–25^. A genome-wide computational study identified a similar motif among those motifs most enriched in human promoters, suggesting a possible functional role ^26^. Our data indicate that the CGCG element is enriched in TATA-less CGI-associated promoters and evolutionarily conserved among vertebrates. Importantly, it is associated with bidirectional transcription only in the context of CGI-associated promoters as assessed by analysis of GRO-Cap and Start-seq datasets that identify sense versus anti-sense nascent transcripts and associated TSS. Using novel reporter constructs, we demonstrate that the CGCG element suffices as a core promoter element to drive bidirectional transcription. Gene Ontology analysis indicates that this element is enriched in the promoters of housekeeping genes, most notably those controlling RNA metabolism and translation, and in promoters producing long non-coding RNAs. Together, our results indicate that the CGCG element functions as a previously unknown driver of CGI-associated TATA-less promoters.

## Results

### Motif discovery in DNase-sensitive CpG islands

Roughly 50 percent of human promoters are associated with a CGI ^27^. To identify novel CGI-associated, independently-functioning promoter elements that potentially drive transcription independent of other promoter elements and are enriched in human CGIs (~30k), we extracted CGI sequences that overlapped with DNase-accessible regions (~192k DNase-seq peaks) in the K562 cell line. We then performed an unbiased motif discovery to identify the most enriched motifs in transcriptionally active CGI-associated promoters (figure 1a). As expected, the SP1 binding site (GC box) had the highest enrichment score consistent with its purported role in driving TATA-less promoters. Binding sites for NRF and ETS were also identified, consistent with roles for these transcription factors in the regulation of CGI-associated housekeeping genes ^28^. We also identified two novel sequence motifs (#7 and #10) that were highly conserved within vertebrates. There were more than 400 incidences of motif #10 that coincided with DNase-seq footprints in multiple cell lines (K562 is shown), suggesting that this motif represents a shared regulatory element (figure 1b, Supplementary figure 1a). Although most CGI-associated promoters contain one copy of the motifs shown in figure 1a, motifs 7 and 10 occur in multiple copies in a given promoter (figure 1c). Genome Ontology and Metagene profile analyses showed that motif 7 and 10 are enriched significantly in annotated human CGI-containing promoters, with motif 10 being far more enriched in promoters of annotated coding and non-coding genes despite being less frequent (figure 1b; motif 7=1408 vs. motif 10=413 copies) (figure 1d).

**Figure 1.**
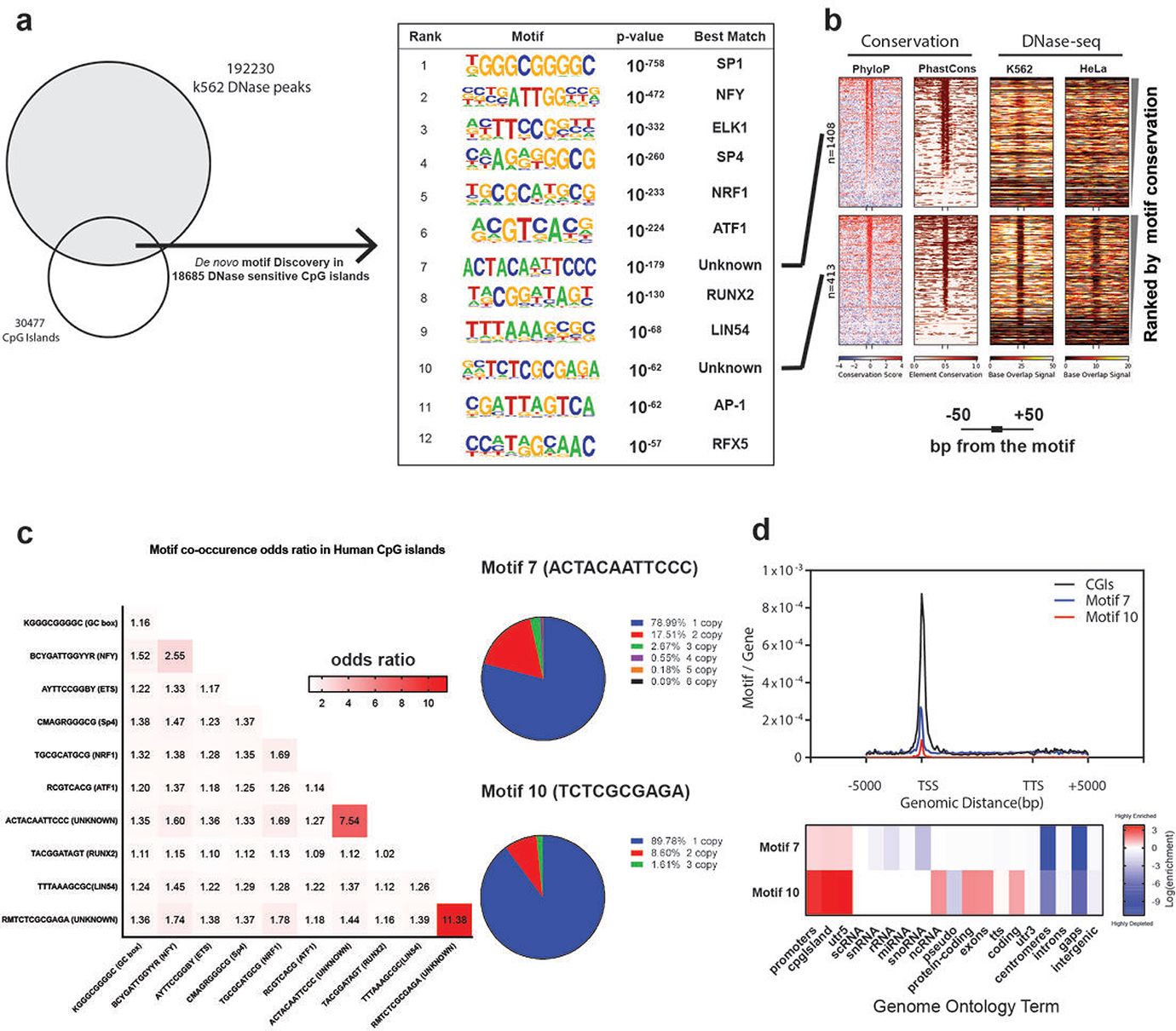
Identification of enriched motifs in human CGIs. a) The intersecting region between K562 DNase-accessible peaks and CpG islands was used to identify enriched regulatory motifs. Among other known transcription factor binding sites, two previously uncharacterized motifs, #7 and #10, were identified. b) Base-wise (PhyloP) and predicted conserved elements (phastCons) score profiles of motif number 7 and 10 in human CGIs and the flanking 50 nucleotides highlight the conservation of these two motifs. Both motifs occur in DNase sensitive CGIs of K562 and other cell lines (Supplement figure 1a). In contrast to motif 7, motif 10 exhibits a marked DNase-seq footprint profile in CGI-associated promoters. c) Motif co-occurrence odds ratio matrix in DNase-sensitive CGIs. The odds ratio is the ratio of observed motif co-occurrence divided by what is expected if motifs were distributed by chance. d) Metagene profile, generated by Homer package, for all CGIs, motif 7 and 10 shown relative to the gene bodies. Annotation enrichment scores in the genome were calculated using the cumulative hypergeometric distribution method found in the Homer package.

### CGCG elements recruit transcriptional machinery and activate gene expression

To determine whether motif 7 and 10 could confer transcriptional activity independently, we cloned the sequence of the most common variant of each motif (ACTACAATTCCC and TCTCGCGAGA, respectively) into the promoterless firefly luciferase reporter construct, Empty pGL2-basic. The resulting constructs were then separately cotransfected along with a control reporter for Renilla luciferase driven by the HSV-1 thymidine kinase promoter (pRL-TK) into human embryonic kidney (HEK293T) cells. Motif 10, but not Motif 7, significantly activated firefly reporter gene expression (figure 2a). This result encouraged us to focus on motif 10, which we named the “CGCG element” based on its central motif. A genome-wide analysis found that this element maps within 50bp of annotated TSSs in human and mouse genomes (Supplementary figure 1b) suggesting that this element could potentially function as a core promoter element ^29^. To address the function of a specific naturally-occurring CGCG element, we analyzed the CGI-containing promoter of the human Density Regulated gene (*DENR*). The *DENR* promoter contains three tandem CGCG elements separated by 21 and 11 nucleotides (figure 2b). To determine the role of each CGCG element in this promoter, we inserted promoter fragments containing CGCG #1, CGCG #1,2 and CGCG #1,2,3 into pGL2-basic. Although a single copy of the CGCG element significantly increased reporter activity, there was a 7-and 17-fold increase in reporter activity with the addition of the second and third CGCG elements, respectively. Introducing G to T mutations in all CGCG elements (CTCG #1,2,3) dramatically decreased promoter activity, suggesting that the CGCG element is necessary and sufficient to drive reporter expression and that there is a cooperativity between multiple CGCG elements (figure 2c).

**Figure 2.**
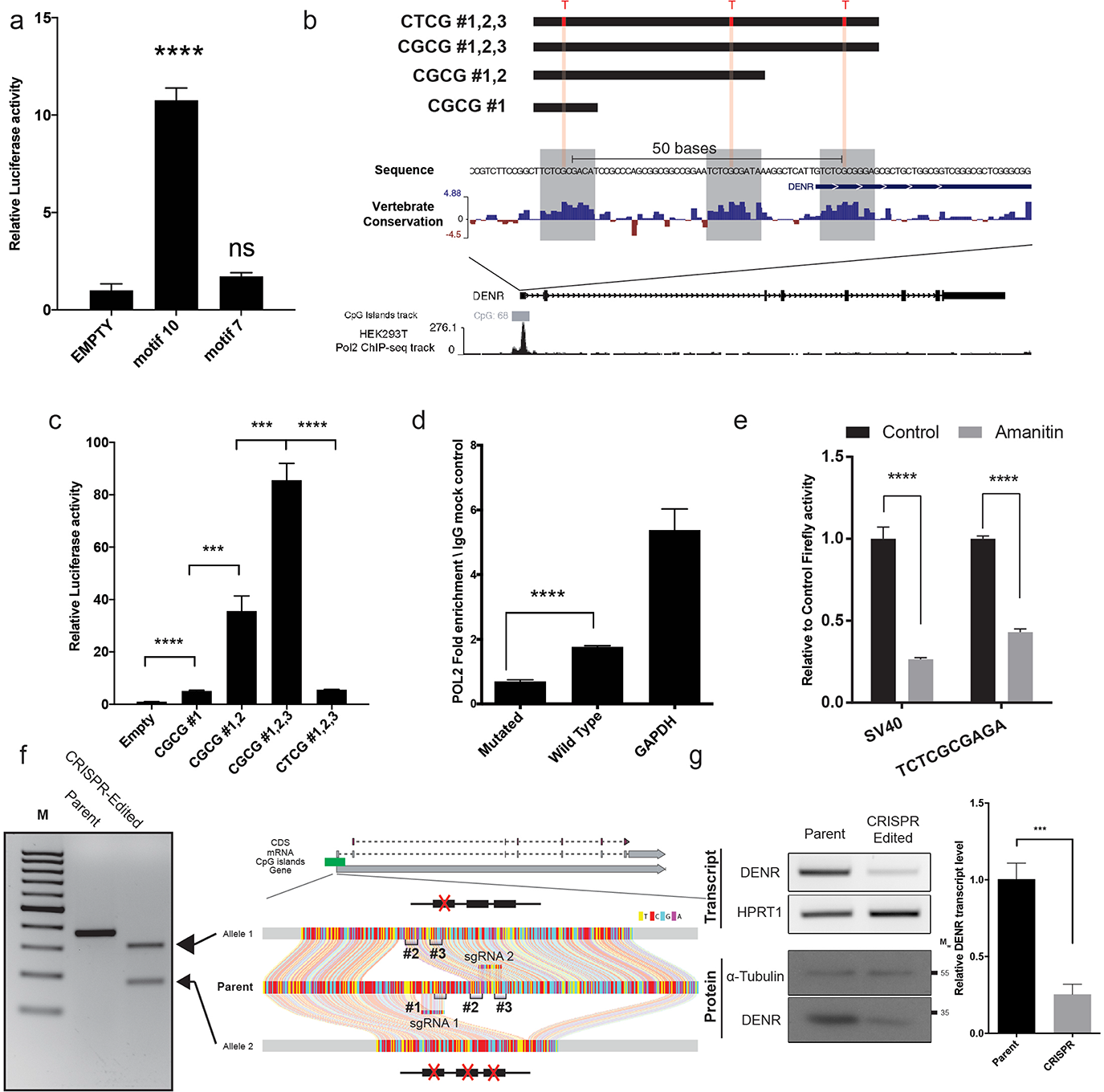
CGCG elements recruit RNA polymerase 2 and activate reporter expression. a) Firefly reporter activity driven by motif 7 and 10. b) The structure of human *DENR* promoter and promoter fragments used for reporter studies. *DENR* promoter encompasses three highly conserved copies of the CGCG elements. The ENCODE POL2 ChIP-seq performed on HEK 293T cells shown in the bottom demarcates the POL2 occupancy in the region. c) Reporter activity of the corresponding *DENR* fragments as described in section b of this figure. d) POL2-ChIP using the wild-type (TCTCGCGAGA) and mutant (TCTC<u>T</u>CGAGA) reporter construct. Human *HPRT* promoter was used as a positive control. e) The effect of α-amanitin on TCTCGCGAGA-driven firefly reporter. f) CRISPR/Cas9 double-nickase strategy to target CGCG elements in the *DENR* promoter. Ribbon plots show Sanger sequences of parental and edited alleles in a clone that contained a microdeletion in the *DENR* promoter. g) The resulting genome editing critically affected DENR transcription as assayed by RT-PCR and quantitative RT-PCR. This reduction in transcription resulted in lower DENR protein levels as determined by immunoblotting. Data are represented as the mean of three replicates ± SD.

To determine if CGCG element-driven gene expression is dependent on RNA polymerase 2 (POL2), we transfected HEK293T cells with reporter constructs that contain either the consensus motif (TCTCGCGAGA) or a CTCG mutation (TCTC<u>T</u>CGAGA) and performed a chromatin immunoprecipitation (ChIP) for POL2 ^30^. As shown in figure 2d, POL2 bound the wild-type (WT) CGCG but not to the mutant C<u>T</u>CG site. Analysis of the POL2 ChIP-seq ENCODE dataset in HEK293T cells identified POL2 binding peaks coincident with CGCG elements in the *DENR* promoter (figure 2b). α-amanitin, a POL2 inhibitor ^31^, decreased CGCG element-driven reporter expression (figure 2e), suggesting that POL2 is indispensable for CGCG dependent gene expression.

To assess the effect of removing CGCG elements on the endogenous *DENR* promoter activity, we employed a CRISPR/Cas9 double-nickase strategy ^32^ to delete a small CGCG-containing *DENR* region in the HEK293T cell line. One cell clone, containing a deletion of approximately 200 base pairs (bp) removed all three CGCG elements in one allele, and a separate 100bp deletion removed one of the CGCG elements in the other allele without affecting the remaining CGI in the promoter (figure 2f). Removal of these CGCG-containing regions caused a significant decrease in the *DENR* transcript and protein levels compared to WT controls (figure 2g). Together with the reporter analyses, these findings suggest that CGCG elements actively recruit transcriptional machinery and promote gene expression in the CGI-associated promoter of *DENR* gene.

### CGCG element confers bidirectional transcription activity

Due to the palindromic nature of the TCTCGCGAGA motif, we wondered whether the CGCG elements could also activate bidirectional transcription. To test this, we developed a novel bidirectional reporter construct (LuBiDi) to measure promoter activity using firefly and Renilla luciferase genes as reporters of directional transcription from a central control motif (figure 3a).

**Figure 3.**
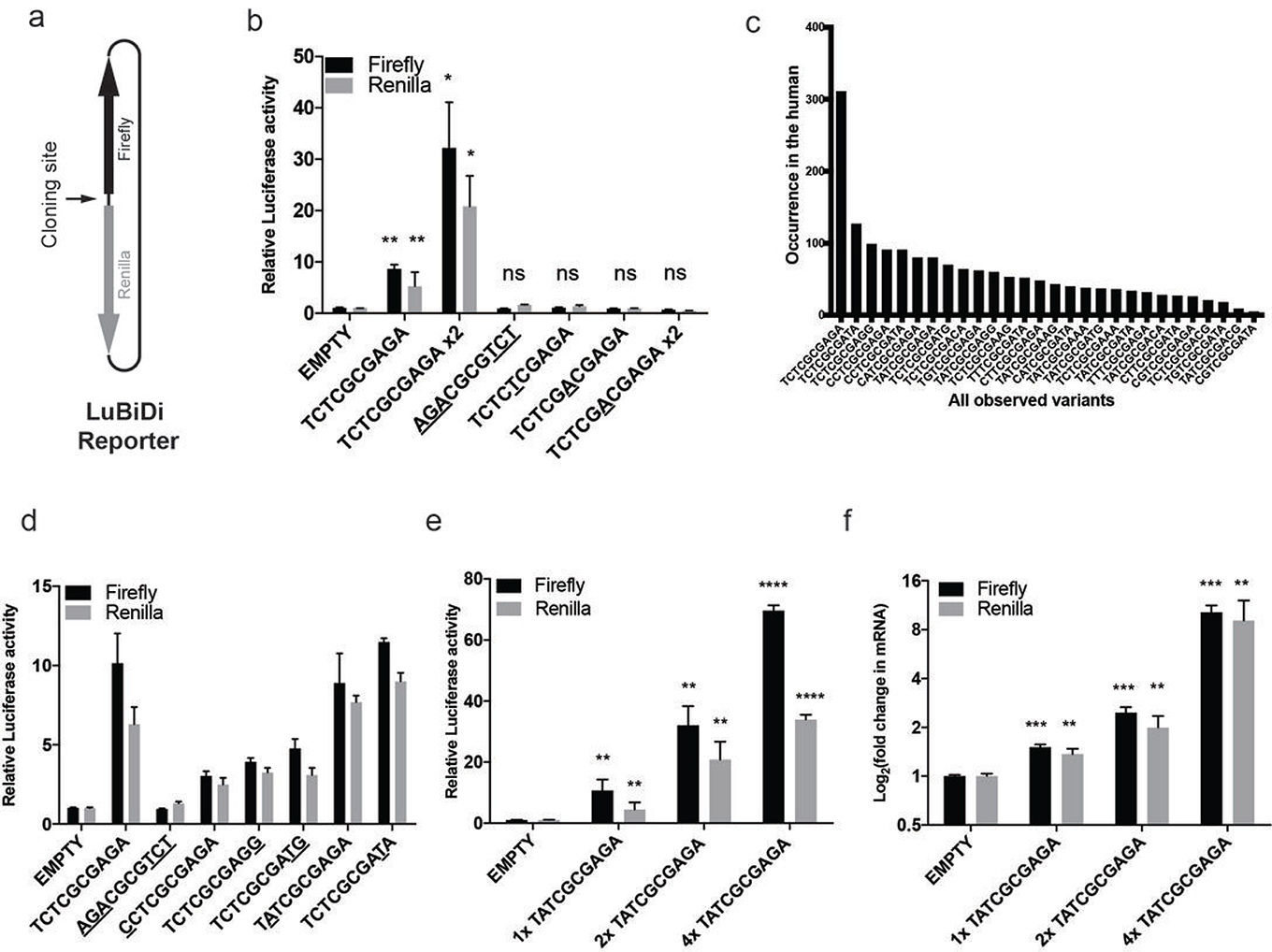
CGCG elements promote bidirectional gene expression in the LuBiDi reporter system. a) The structure of the LuBiDi reporter construct and the cloning site. b) A copy of the TCTCGCGAGA motif inserted in the LuBiDi construct was sufficient to activate the expression of both firefly and Renilla reporters. Flank-exchanged (<u>AGA</u>CGCG<u>TCT</u>), G to T transversion mutation (TCTC<u>T</u>CGAGA) and an insertion mutation in the middle of CGCG (TCTCG<u>A</u>CGAGA) abolished the dual activation. c) The frequency of common CGCG element sequence variants observed in the human genome. d) The bidirectional promoter activity of a few CGCG element naturally-occurring variants. e) Promoter activities of reporter genes were associated with the copy number of T<u>A</u>TCGCGAGA motif present in the LuBiDi. f) Corresponding transcript levels from reporters in section “e” of this figure.

We inserted one or two copies of the TCTCGCGAGA motif into the LuBiDi plasmid and measured reporter activity. A single CGCG element was sufficient to induce both firefly and Renilla reporters whereas two CGCG elements induced an additional 4-fold increase (figure 3b). To study the motif sequence requirement for this activation, we introduced mutations in the motif that disrupted the wild-type sequence in various locations. First, to determine whether the palindromic structure was more important than sequence content in conferring the bidirectional transcriptional activity, we exchanged the flanking sequences to form <u>AGA</u>CGCG<u>TCT</u>, which maintains both symmetry and CpG content. This mutation abrogated the dual activation of reporters (figure 3b), suggesting that the CGCG element has sequence polarity. A CGCG -> CTCG transition mutation (TCTC<u>T</u>CGAGA, reduced CG content) and an “A” insertion into CGCG (TCTCG<u>A</u>CGAGA, unchanged CpG content) abrogated dual reporter activity (figure 3b). The inclusion of two copies of the A insertion mutant failed to induce transcription. Altogether, these results indicate that the WT element, CGCG core plus the flanking palindromic sequences found in motif 10, are required for promotion of bidirectional transcriptional activity.

To analyze the expression dynamics of CGCG elements in single cells, we developed another promoter-less bidirectional reporter (pmCGFP) that codes for enhanced Green Fluorescent Protein (eGFP) and mCherry reporters in opposite directions (Supplement figure 2a). One or three copies of TCTCGCGAGA motifs were inserted into this reporter construct, which were then cotransfected into HEK293T cells along with a CMV promoter construct driving the Blue Fluorescent Protein (BFP) as a transfection control. Cells simultaneously expressed both GFP and mCherry reporter genes starting 12 hours after transfection only for constructs containing the TCTCGCGAGA element (supplement figure 2b). Immunoblot analysis indicated that GFP and mCherry protein levels were proportional to the number of inserted TCTCGCGAGA motifs (Supplement figure 2c). We also tracked individual cells using live imaging microscopy and observed that the two reporter genes are expressed simultaneously after transfection (Supplement figure 2d; Supplementary Video). We also performed a similar imaging experiment using an mCherry reporter fused to the H2b in HEK293T and NMuMG mouse mammary cell lines, again showing simultaneous expression of both reporters (Supplementary Figure 2e, f). Collectively, these results suggest that this element is a potent bidirectional transcription activator in multiple species.

An analysis of human CGI-associated promoters indicated that CGCG elements could also contain less frequent, single nucleotide variations in TCT or AGA flanking sequences (figure 3c). To determine the impact of these minor variations on bidirectional transcription activity, we compared LuBiDi constructs with one TCTCGCGAGA motif to those containing naturally variant sequences, using the AGA <-> TCT flank-exchanged mutant as a negative control (figure 3d, the variation in a specific nucleotide is underlined). <u>C</u>CT, AG<u>G</u> or A<u>TG</u> flanking sequences (underline represents changes) decreased relative dual reporter activity whereas variants that contain A<u>T</u>A or T<u>A</u>T showed similar activity to that of the TCTCGCGAGA motif (figure 3d). The data suggest that some, but not all, variability in the flanking sequences confer core promoter activity, albeit at lower efficiencies compared to the TCTCGCGAGA motif. The data also showed that imperfect palindrome elements can still drive bidirectional transcription.

To study the role of copy number variation on bidirectional transcription activity in more detail, we generated LuBiDi reporters that contain one, two or four copies of T<u>A</u>TCGCGAGA, a common variant of the CGCG element with an imperfect palindrome. Reporter activity increased proportionally with the number of motifs as measured by luciferase activity or luciferase transcript levels (figure 3e, f).

### Endogenous CGCG elements confer transcriptional activity in CGI-associated promoters and methylation abrogates its promoter activity

To determine if CGCG elements are associated with bidirectional transcription from endogenous promoters, we analyzed a previously published GRO-cap (global run-on sequencing followed by enrichment for 5’-cap structure) analysis performed on K562 cells ^33^. GRO-cap allows for the detection of nascent, often unstable strand-specific RNA transcripts that are usually undetectable by common RNA-seq methods, likely because of the greatly increased sequencing depth near to TSS associated with directional transcription of coding RNAs. We found that the bidirectional transcription is associated almost exclusively with CGCG elements that occur in CGI-enriched promoters (figure 4a). Gene Ontology (GO) analysis showed that genes containing CGCG promoter element produce protein-coding transcripts whose products form discernable protein-protein interacting networks (Supplementary figure 3). Specifically, these genes encode core components of RNA metabolism and the translational apparatus (Table 1).

**Figure 4.**
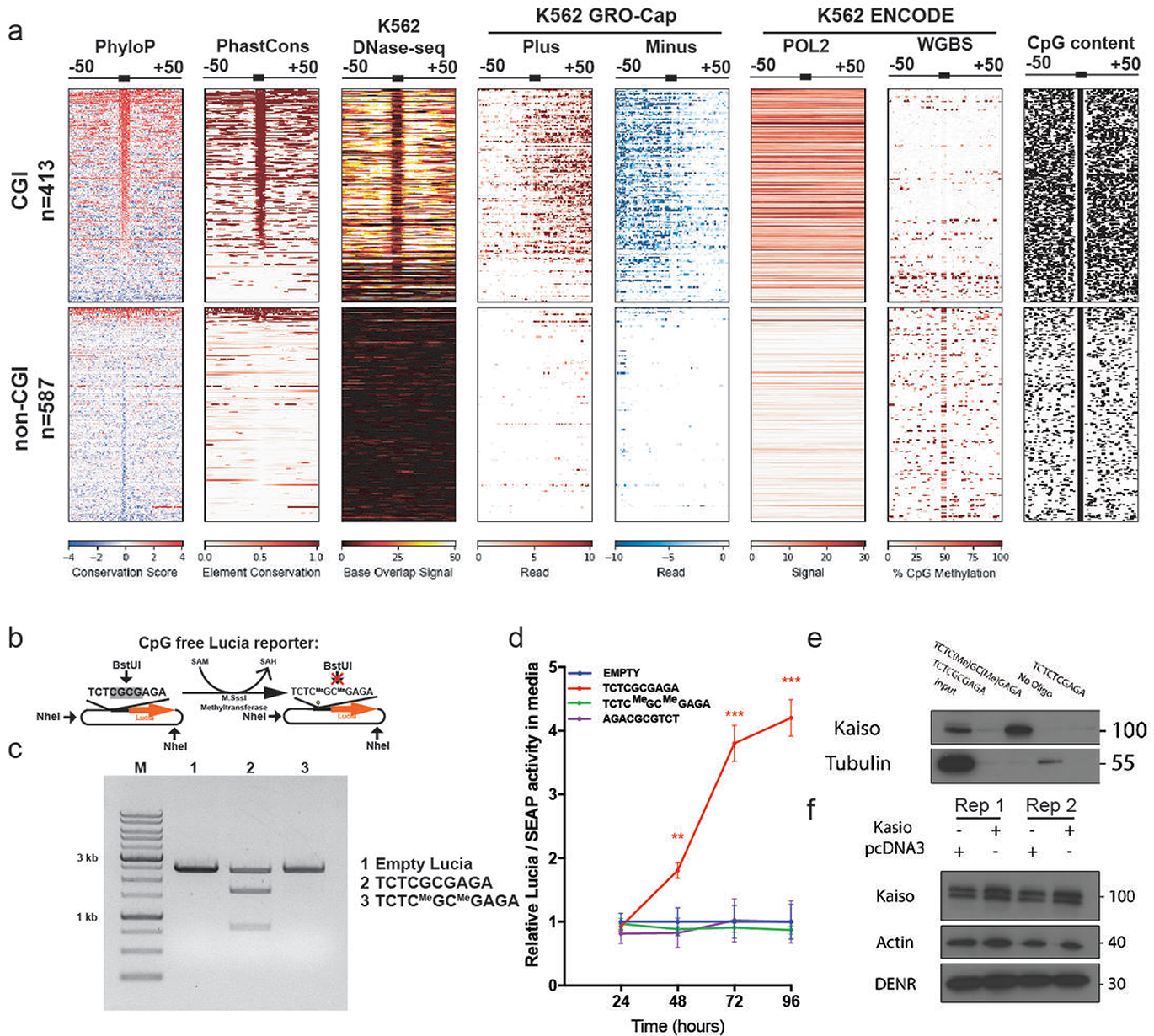
CGCG elements are transcriptionally active in CpG islands and methylation abolishes its activity. a) CGCG elements that occur in CGIs mark DNase-seq footprint in DNase-sensitive regions and associated with divergent plus and minus GRO-Cap transcripts in K562 cell line. The occupancy of POL2 on CGCG elements in CGIs as gauged by ENCODE POL2 ChIP-seq performed in K562 cell line. ENCODE WGBS methylation data for K562 cell line showed the percentage of CpG methylation in CGIs and non-CGI sites. b) TCTCGCGAGA was inserted into a CpG-free Lucia reporter construct. The construct was methylated using M.SssI CpG methyltransferase and SAM as the methyl donor. c) Methylation of TCTCGCGAGA in the construct assessed by agarose gel analyses after digestion with NheI and BstUI enzymes. BstUI restriction enzyme recognizes nonmethyl CG/CG sequence and performs a blunt cut (/ indicates the BstUI blunt cut site). d) The reporter construct containing methylated TCTCGCGAGA did not activate Lucia activity. Data are represented as the mean of three replicates ± SD. e) Oligonucleotide pull-down followed by immunoblotting for Kaiso protein. f) Ectopic transient (72h) overexpression of Kaiso protein in HEK293T cells did not alter DENR protein levels.

**Table 1.** Top GO terms for genes whose promoters contain CGCG elements. FDR: False Discovery Rate. Please see methods section for the definition of N, B, n and b variables.

| Term | P-value | FDR | Enrichment (N, B, n, b) |
| --- | --- | --- | --- |
| nuclear-transcribed mRNA catabolic process, nonsense-mediated decay | 1.21E-08 | 1.71E-04 | 5.45 (13906,109,398,17) |
| Co-translational protein targeting to membrane | 2.57E-08 | 1.82E-04 | 5.96 (13906,88,398,15) |
| mRNA metabolic process | 1.97E-07 | 9.26E-04 | 2.43 (13906,576,398,40) |
| amide biosynthetic process | 2.77E-07 | 9.80E-04 | 3.31 (13906,253,398,24) |
| protein targeting to ER | 3.26E-07 | 9.21E-04 | 5.32 (13906,92,398,14) |
| establishment of protein localization to endoplasmic reticulum | 4.89E-07 | 1.15E-03 | 5.15 (13906,95,398,14) |
| protein targeting to membrane | 5.69E-07 | 1.15E-03 | 4.21 (13906,141,398,17) |
| SRP-dependent co-translational protein targeting to membrane | 6.11E-07 | 1.08E-03 | 5.47 (13906,83,398,13) |
| protein localization to endoplasmic reticulum | 1.34E-06 | 2.10E-03 | 4.75 (13906,103,398,14) |
| nuclear-transcribed mRNA catabolic process | 3.01E-06 | 4.25E-03 | 3.57 (13906,176,398,18) |

CpG dinucleotides in CGI-associated promoters are invariably unmethylated ^13^, we asked if the methylation state of the CGCG elements might explain the observation that only the elements within CGIs are transcriptionally active. Analysis of ENCODE Whole Genome Bisulfate Sequencing (WGBS) from K562 cells indicated that in contrast to CpG-poor regions of the genome, CGCG elements in CGIs are largely unmethylated (figure 4a). This observation prompted us to determine experimentally whether CpG methylation could alter the promoter activity of the CGCG element. We cloned a single copy of TCTCGCGAGA into a secretory luciferase reporter construct that is devoid of CpG sequences (CpG-free Lucia). In this construct, the only CpG sequences are the ones contributed by the CGCG element (figure 4b). The CpG sequences were then fully methylated using M.SssI CpG methyltransferase, confirmed by saturated methyl-sensitive enzymatic digestion (figure 4c). In comparison to the high reporter activity induced by the unmethylated TCTCGCGAGA-containing construct, methylation abrogated the promoter activity (figure 4d), strongly suggesting that CGCG methylation antagonizes it promoter function.

A transcription factor zBTB33, also known as Kaiso, was shown previously to be enriched on methylated “CGCG” nucleotides ^34^. Kaiso has been shown to interact with the repressive complex SMRT, leading to suppression of gene expression ^35^. As illustrated in figure 4e, this transcription factor interacts only with the methylated CGCG element confirming previous observations ^36^. The transient overexpression of Kaiso in HEK293T cells did not significantly alter the endogenous DENR protein level (figure 4f). These results indicate that Kaiso does not bind to the CGCG element when it is not methylated. Since Kaiso does not suppress the DENR promoter activity when expressed in 293T cells, it is suggested that the CGCG element in the DENR promoter is not methylated in vivo. Thus, Kaiso along with other zBTB family members likely suppress the CGCG element-driven gene expression only when this element is methylated.

### The CGCG element activates gene expression in different promoter configurations

Given that the CGCG element drives bidirectional transcription, we were interested to determine the frequency of this element in annotated uni-vs. bidirectional promoters. The vast majority of CGCG elements (93%) occur in annotated unidirectional promoters that drive coding or lncRNAs, while 7% occur in an annotated bidirectional promoter (Table 2). However, recent studies suggest that the majority of what were classically defined as unidirectional promoters produce unstable “promoter upstream transcripts” (PROMPTS) ^37^. Based on this, we investigated the role of CGCG elements in three different endogenous promoters that differ in their annotated directionality and whether they combine CGCG element with TATA-boxes. In order to determine the role of endogenous CGCG elements, we simultaneously disrupt CGCG element but maintained CG content by exchanging the flanking sequences (i.e. TCTCGCGAGA to <u>AGA</u>CGCG<u>TCT</u>). We first focused on the *POLR1C/YIPF3* bidirectional promoter region, which has two TSS separated by 30 nucleotides that flank a single CGCG element. We inserted a promoter fragment (~30bp) containing the wild-type CGCG element into the LuBiDi construct, and as a comparison, constructs were generated in which the flanking sequences (AGA and TCT) were exchanged. The WT fragment from *POLR1C/YIPF3* promoter induced bidirectional expression irrespective of its orientation (figure 5a). In contrast, the flank-exchanged mutants, regardless of insert orientation, did not show any discernable reporter activity.

**Table 2.** Frequency of the CGCG elements in annotated promoters.

| Annotated configuration | CGCG elements | Percent |
| --- | --- | --- |
| Unidirectional coding | 364 | 80 |
| Bidirectional coding pair | 22 | 5 |
| Unidirectional non-coding | 58 | 13 |
| Non-coding and coding pair | 9 | 2 |

**Figure 5.**
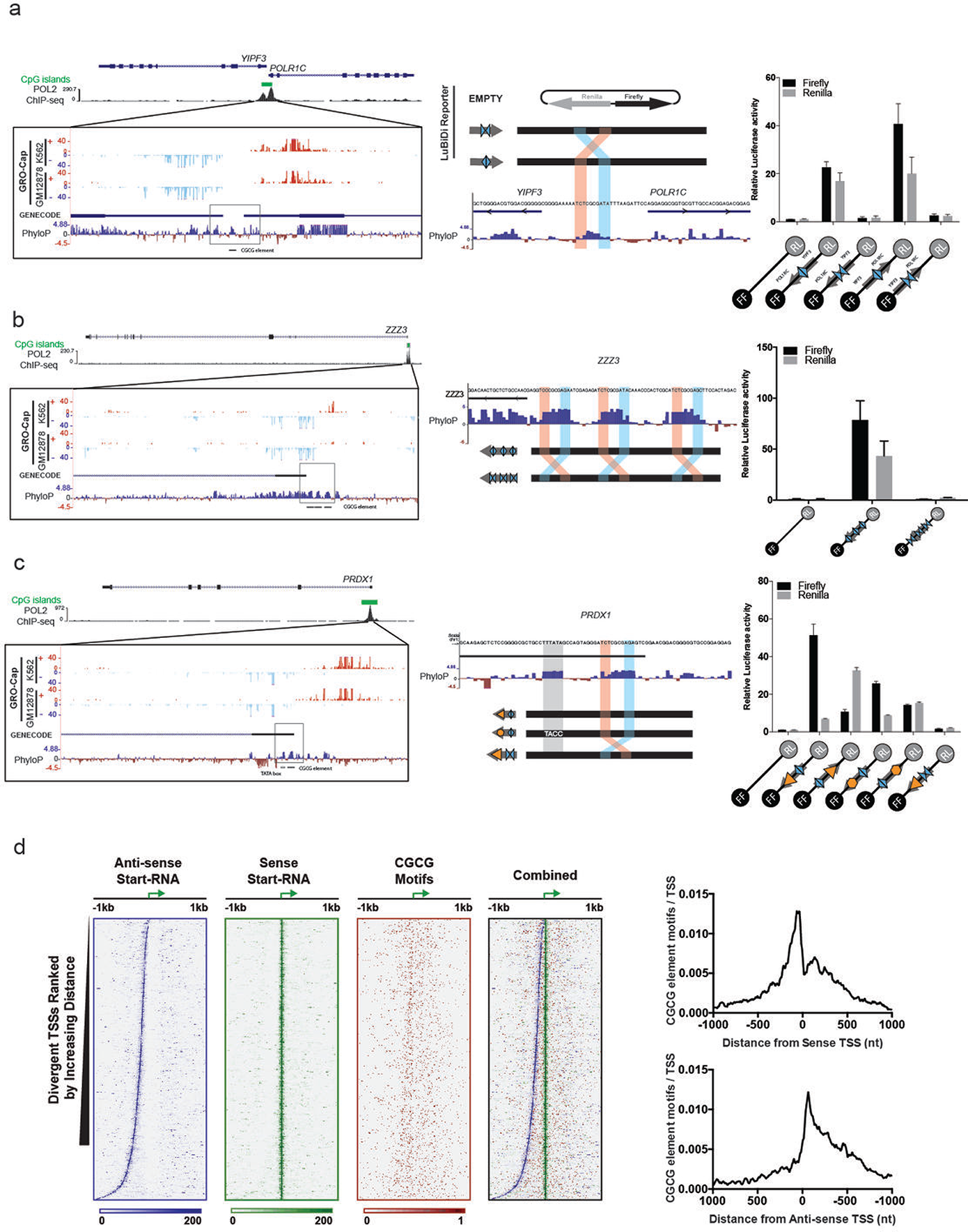
CGCG element in CGI promoters drives gene expression. a) The bidirectional promoter of *POLR1C/YIPF3* gene pairs contains a conserved CGCG element between annotated TSSs. b) The *ZZZ3* promoter contains three copies of CGCG elements. Although this promoter is annotated as unidirectional, the GRO-Cap analysis indicated associated divergent transcripts on the opposite strand. Wild-type fragment of this promoter that contains these three elements, but not the flank-exchanged mutants, confer bidirectional activation of reporter genes. c) The promoter of *PRDX1* gene contains both TATA-box and a CGCG element. GRO-Cap signals show a major TSS 26 nucleotides downstream of the TATA box. Promoter activity associated with this promoter structure indicated that increased directional promoter activity depended on the arrangement of the TATA box. Disruption of TATA-box (CCTA) attenuated this directional activity and the flank-exchanged mutant of the CGCG element abrogated the reporter activity. d) Start-seq data analysis of CGCG elements in the mouse genome. CGCG elements occur mostly within 50bp of sense and anti-sense Start-seq TSSs. Data are represented as the mean of three replicates ± SD.

Next, we analyzed the *ZZZ3* promoter which is similar to the *DENR* promoter in that it contains three CGCG elements (figure 5b). Although the promoter is annotated as directional, PROMPTs on the opposite strand in both the K562 and GM12878 GRO-Cap datasets were found (figure 5b, UCSC genome browser plot). To determine whether these elements are responsible for the *ZZZ3* divergent transcript, we inserted CGCG elements or flank-exchanged elements, from the *ZZZ3* promoter into a LuBiDi construct. As shown in figure 5b, WT sequences but not flank-exchanged could induce bidirectional reporter expression. An analysis of the *DENR* promoter also showed that their three CGCG elements drive bidirectional transcription in LuBiDi and disruption of CGCG core sequences with A insertions abrogated the bidirectional promoter activity (Supplementary figure 4).

We also studied the *PRDX1* promoter, a rare example in which both a single CGCG element plus a TATA-box map within the CpG enriched promoter ^38^. An analysis of GRO-Cap datasets indicated a predominant TSS approximately 25 nucleotides downstream of the TATA-box (figure 5c), yet divergent transcripts were found starting roughly 50-70 bp upstream of the coding region in both K562 and GM12878 cells. To investigate the role of the TATA-box in this configuration, we inserted a fragment containing the TATA-box and CGCG element from this promoter into LuBiDi. We also produced mutants including one that disrupted the first TA in the <u>TA</u>TA-box with CC sequences and another in which the TATA-box orientation was reversed relative to the CGCG element. The WT *PRDX1* promoter mainly drove unidirectional downstream transcription (figure 5c) although some opposite direction reporter activity was noted.

Mutation of the TATA-box severely attenuated downstream directional promoter activity (figure 5c). Interestingly, the reporter containing a flank-exchanged CGCG element did not show any reporter activity even in the presence of a WT TATA-box, suggesting that the CGCG element not only promotes divergent transcription but also acts as a required activator for the TATA-box in this promoter.

To further study the role of CGCG elements in the context of bidirectional promoters, we analyzed a set of mouse bidirectional promoters previously defined using Start-seq ^39^ We assessed the presence of CGCG elements throughout the intervening region in such bidirectional promoters. The coupled sense/anti-sense TSS form boundaries that flank a nucleosome-depleted region (NDR), characterized by an open chromatin structure that permits high accessibility for transcriptional machinery (figure 5d). This analysis indicated that although CGCG elements do not show a fixed distance to sense or anti-sense Start-seq TSSs, they are found mostly in NDRs of mouse bidirectional promoters.

### CGCG elements promote transcription through divergent TSS

Previously identified core promoter elements such as the TATA box and the TCT motif promote transcription through a focused putative TSS that occurs either at a fixed distance downstream (in the case of TATA box) or on a specific nucleotide within the element in the case of TCT motif ^40^. To map the bidirectional TSSs associated with the CGCG element, we employed 5´ RACE (5´ Rapid Amplification of cDNA Ends) using RNA extracted from HEK293T cells transfected with LuBiDi reporter constructs along with pEGFP as a transfection control (figure 6a). This robust method has been successfully used to determine the TSS of many genes in human and other organisms previously ^41, 42^. As shown in Figure 6b, 5´ RACE produced major single products for firefly and Renilla transcripts from a LuBiDi construct containing one copy of the TCTCGCGAGA motif. Sequencing of the resulting RACE products showed a preference for A or G as the +1 nucleotide, and C or T as the -1 nucleotide, conforming to the previous observation that ideal TSS tend to use pyrimidines and purines at the -1 and +1 positions, respectively ^38^. Although multiple TSSs were found in the sense or anti-sense directions, there was a predominant firefly TSS (7 of 25 clones) 28 nucleotides and a predominant Renilla TSS (9 of 21 clones) 51 nucleotide from the TCTCGCGAGA element (figure 6c). However, a majority of preferred Renilla TSS were downstream of the Renilla initiation codon (ATG), and thus, unlikely to produce active Renilla luciferase product. This likely explains why the relative Renilla luciferase activity is always lower than that of the firefly as was previously observed in figure 3b.

**Figure 6.**
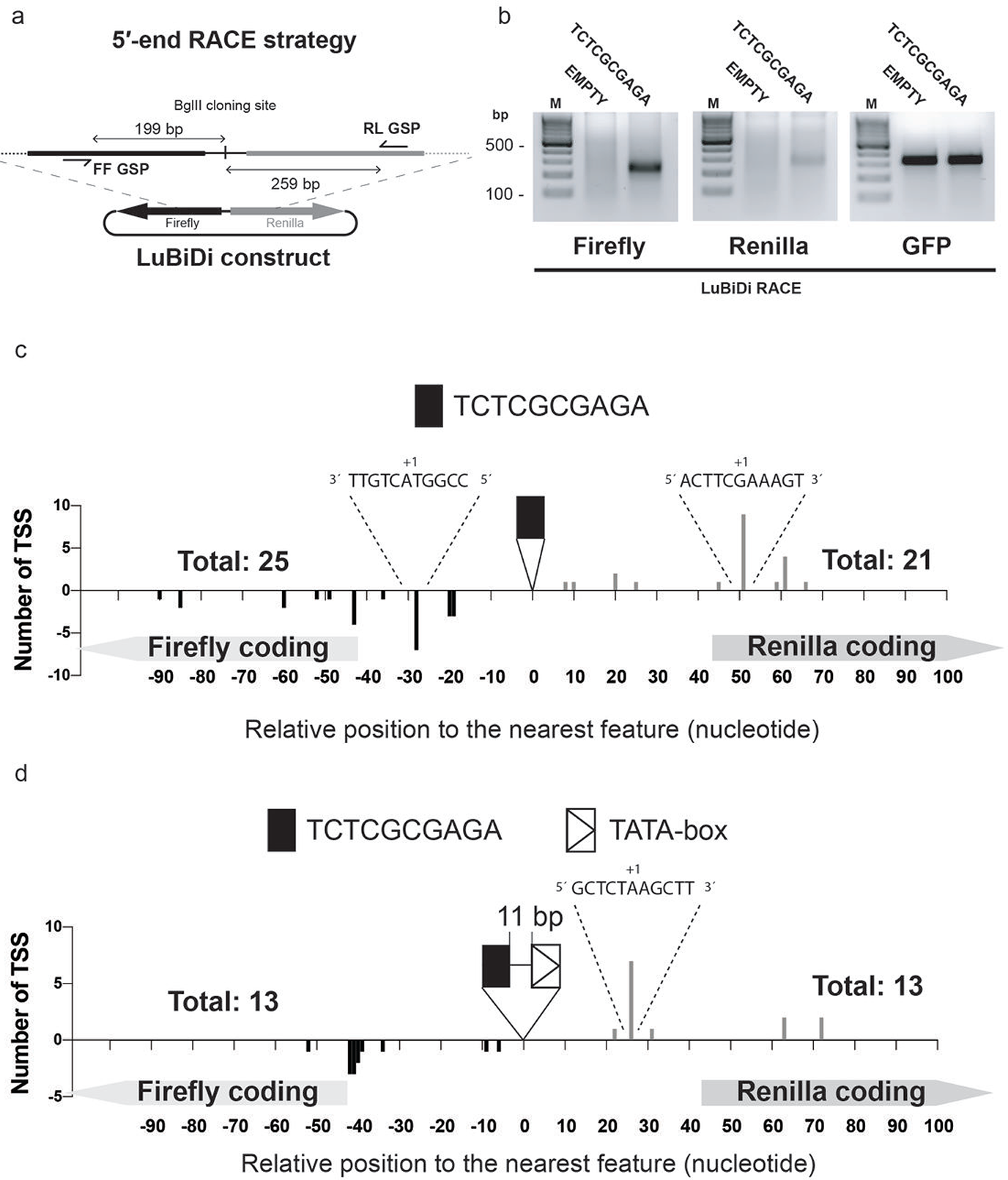
CGCG element is associated with bidirectional transcription start sites. a) The location of gene-specific primers (GSP) used in our 5´ RACE experiments to identify bidirectional TSSs in the LuBiDi based reporter constructs. Firefly and Renilla primers were designed 199 and 259bp away from the Bglll cloning site, respectively. b) Agarose gel image of firefly and Renilla RACE PCR products for the LuBiDi constructs containing none or one copy of TCTCGCGAGA. GFP transcript was used as an internal control for the RACE experiment. c) TSS were determined for the LuBiDi construct containing a copy of the TCTCGCGAGA motif. TSS locations are indicated in nucleotide relative to the CGCG element. The sequence of +1 nucleotide and flanking five nucleotides are also shown on major TSSs. d) Divergent TSS for the LuBiDi construct that contained a TATA-box and a CGCG element from the *PRDX1* promoter. TSS positions are indicated in nucleotide relative to the nearest feature (the CGCG element or the TATA-box). Note that positive TSS counts were used for the Renilla transcripts and negative numbers for the firefly transcripts. The number of sequenced clones for each reporter constructs are indicated above coding regions.

Next, we determined how the presence of a TATA-box affects CGCG element-driven transcription from the LuBiDi reporter containing both elements from the *PRDX1* promoter. In this construct, the TATA-box is arranged between the Renilla reporter and the CGCG element. Sequencing of the Renilla RACE products showed a predominant TSS (7 of 13 clones) 26 nucleotides downstream of the TATA box on the Renilla-coding strand (figure 6d). In contrast, on the firefly reporter coding strand, there was a concentration of multiple TSSs 40-43 nucleotides downstream of CGCG element. This TSS pattern differs from those induced from the construct containing one copy of the CGCG element (figure 6c). Together with reporter data presented in figure 5c, these results suggest that the CGCG element and TATA box cooperate to induce transcription in the *PRDX1* promoter.

## Discussion

In this study, we identify a novel promoter element that drives bidirectional transcription mainly in the context of TATA-less promoters. Whereas other promoter elements (e.g. TATA and GC boxes) require an activator binding site to initiate directional transcription ^6^, a single instance of the CGCG element is both necessary and sufficient to promote bidirectional transcription. However, in comparison to other known core promoter elements, which typically occur once in most promoters, CGCG elements occur in multiple copies in small percentages of CGI-containing promoters, a phenomenon that could potentially dictate RNA polymerase recruitment and consequent transcriptional rates.

An interesting yet poorly studied feature of vertebrate genomes is the presence of CpG rich regions known as CGIs ^14^. Although CGIs mark transcriptionally active regions of the genome, the mechanism of RNA polymerase recruitment in these regions has been elusive ^13^. Through enrichment analysis, we found that CGCG elements are enriched in CGI-containing promoters and that they can recruit transcriptional machinery to promote bidirectional transcription, a feature that most transcriptionally active CpG islands was shown to possess ^19^. Additionally, we provide evidence that in some rare cases, the CGCG element could interact functionally with an adjacent TATA-box within a CGI to activate gene expression. Similar synergetic activities have been described previously ^43, 44^ suggesting that the CGCG element also shares this attribute with other known core promoter elements.

How housekeeping genes whose products are core components of cellular processes are transcriptionally regulated is poorly understood. In this study, we found that genes whose products play a central role in translation and transcription are enriched for CGCG elements in their CGI-associated promoters. This analysis led us to identify a group of ribosomal genes whose CpG rich promoters contain one or multiple copies of CGCG elements (Supplementary Figure 5). These promoters do not contain the previously described TCT motif that is thought to regulate the transcription of the other group of ribosomal genes in humans ^40^. These results suggest that TCT and CGCG elements regulate the expression of different sets of ribosomal genes in human. In addition to genes encoding ribosomal proteins, promoters of key translation initiation factor genes encoding EIF5, EIF3H, and DENR, as well as the essential translation termination factor ETF1, contain copies of the CGCG elements. This is consistent with the current perspective that different classes of promoter elements regulate functionally distinct protein coding genes ^1^.

Additionally, we directly demonstrated that methylation of CpGs in the CGCG element could suppress its promoter activity. Indeed, roughly 80 percent of CpG sites in the genome, particularly CpGs that occur outside of CGIs, are methylated ^45^. We speculate a switch-like mechanism that could activate or repress gene expression based on the methylation status of CGCG elements. Accordingly, we propose a model where CGCG elements, when occurring in CGIs, are protected from methylation thereby maintaining promoter activity in housekeeping genes. In contrast, CGCG elements in other regions of the genome would be more subject to methylation, resulting in transcriptional silencing. In theory, DNA methylation of CGCG elements could protect the genome from spurious transcription, as reviewed elsewhere ^46^. A similar switch-like mechanism for a group of transcription factors that contain CpG motif has been described in the past in which CpG methylation would affect the affinity of transcription factors such as Kaiso ^47^. Although the nature of the factor, or factors, that bind to non-methyl CGCG element has yet to be clarified, our results suggest that ChIP-seq studies should be interpreted with greater consideration to account for the differential binding of proteins to methyl or non-methyl CpG-containing motif sequences.

In a recent study, Dual Specificity Kinase 1 (DYRK1A) was identified as a novel POL2 C-terminal domain (CTD) kinase and activator of RNA polymerase 2 ^48^. Subsequent ChIP-seq analysis of DYRK1A showed that this protein is specifically enriched in CGCG containing promoters. It has been suggested that RNA polymerases are recruited through various transcriptional preinitiation complexes (PIC) that specifically regulate different promoter classes ^1, 49^. Therefore, we speculate that CGCG elements directly or indirectly recruit DYRK1A as the component of a novel PIC that remains to be completely elucidated.

In conclusion, this study provides strong evidence that the CGCG element is evolutionarily conserved in vertebrates, functioning as an active component of CGI-associated promoters. The unmethylated form of the element may be sufficient to drive bidirectional transcription of TATA-less promoters. With the identification of the CGCG element interacting factor or factors in the future, we may soon gain a better picture of how basal transcription of TATA-less housekeeping genes is regulated.

## Materials and Methods

### Cell culture and treatments

Human embryonic kidney 293T and NMuMG cell line were cultured in Dulbecco’s Modified Eagle Medium (DMEM) media supplemented with 10% fetal bovine serum, penicillin and streptomycin antibiotics. Cell lines were grown in an incubator at 37°C and 5% CO_2_.

For the α-amanitin treatment experiment, HEK293T Cells were transfected with SV40 promoter-driven firefly reporter (pGL2-pro), or a construct containing a copy of TCTCGCGAGA. 24 h post-transfection, cells were treated with 5 μg/ml α-amanitin (Santa Cruz) as described ^50^ or with PBS (control), and firefly and Renilla luciferase bioluminescence activities were measured 24h after treatment.

### Reporter constructions and assays

One to three copies of the CGCG elements from *DENR* promoter were synthesized as double stranded oligonucleotides (IDT DNA) and cloned into the BglII and MluI restriction sites of a luciferase reporter construct that lacks promoter sequences (pGL2-basic, Promega). 1 μg of cloned reporter DNA along with 100 ng of a Renilla reporter construct (pRL-TK) as transfection control were transfected into HEK293T using Roche X-tremeGENE 9 (Roche) transfection reagent according to manufacturer’s protocol. The luciferase activities were measured 24 h after transfection according to the Dual Luciferase assay protocol (Promega).

Luciferase bidirectional (Empty-LuBiDi) reporter was constructed by PCR amplification and subsequent cloning of the firefly luciferase gene from pGL2-Basic into the BglII site of promoterless Renilla cassette from the pRL-Null plasmid and followed by site-directed mutagenesis to remove secondary BglII recognition site downstream of firefly poly-A site. The primer sequences used are available in supplementary information 1. Bioluminescence assays were performed as described above except that transfection was normalized by co-transfecting with a vector that expresses secretory alkaline phosphatase (pSELECT-zeo-SEAP, Invivogene) into the medium.

For the construction of the bidirectional fluorescence reporter, pmCGFP, we PCR amplified and cloned the h2b-mCherry fused gene (plasmid Addgene id #20972) head-to-head into a promoterless eGFP containing construct. The resulting construct (eGFP + h2b-mCherry) was then digested with AgeI to release h2b-coding fragment and auto-ligated to generate the pmCGFP (eGFP + mCherry). Double strand oligonucleotides encoding one or three copies of TCTCGCGAGA into the AgeI restriction site of this reporter.

For CpG free reporter and methylation experiments, an oligonucleotide encoding single copy of TCTCGCGAGA was inserted into HindIII restriction site of pCpGfree-basic-Lucia (Invivogen). 10 μg of purified plasmid was incubated with 10 enzymatic unit (U) M.SssI methyltransferase (NEB) supplemented with fresh 100 μM S-adenosyl methionine (SAM) as the methyl donor in 37°C for 8h. DNA was extracted using phenol-chloroform followed by ethanol precipitation. The DNA was incubated for another 8h after addition of 10 U M.SssI which was followed by DNA extraction as described before. As a control, a mock reaction was also carried out lacking M.SssI enzyme. To test the methylation efficiency, we digested 300ng of reporter constructs using 10 U of NheI and BstUI for 30 min. Because CGCG methylation blocks BstUI cleavage, EMPTY and methylated construct digested only by NheI enzyme producing two indistinguishable bands at 2.4 kb. However, unmethylated TCTCGCGAGA which is cut by BstUI enzyme, as well as NheI, produced three smaller bands The sequences of inserts for each promoter fragment and related mutations are provided in the Supplementary information.

### qRT-PCR

HEK 293T cells were transfected with 1 μg of LuBiDi reporters containing 0, 1, 2, 4 copies of TCTCGCGATA. Cells were lysed after 24 h using TRIzol (Life Technologies), and RNA was extracted using a chloroform-isopropanol protocol. RNase-free DNase I (Thermo Fischer Scientific) was then used to digest contaminating DNA followed by extraction by acidic-phenol chloroform protocol, precipitated using ethanol and dissolved in RNase-free water. 1 μg of the resulting purified RNA was used to prepare cDNA using M-MLV reverse transcriptase according to manufacturer’s recommended protocol (Life Technologies). Transcript levels were measured using iTaq Universal SYBR Green Supermix (Bio-Rad) on an ABI-7900 RT-PCR instrument. Transcript levels were normalized using primers for *HPRT1*. Primers designed to amplify the bacterially expressed AMP resistant gene in the LuBiDi construct were used as negative control to rule out plasmid contamination. Melting curves analyses for all PCR experiments were performed to validate faithful amplification of PCR products. Information on primer sequences is described in Supplementary Information 1.

### Chromatin Immunoprecipitation (ChIP)

ChIP was performed according to a protocol described in Lee, et al. ^51^. In brief, 10 million HEK 293T cells were cultured in 15 cm dishes and transfected with 10 μg reporter DNA using X-tremeGENE 9 transfection reagent. 48 h post-transfection cells were treated with the cross-linking reagent formaldehyde (1% in PBS, Sigma) for 5min. Glycine solution (0.125 M) for 10min was used to stop the cross-linking reaction. Followed by 2 washes with ice-cold PBS. 10 million cells were lysed with lysing buffer (50 mM HEPES-KOH, pH 7.5, 140 mM NaCl, 1 mM EDTA, 10% glycerol, 0.5% NP-40, 0. 25% Triton X-100, 1X protease inhibitors), and their nuclei were isolated by centrifugation (5 min, 1000 RPM) and then sonicated in sonication buffer (10 mM Tris-HCl, pH 8.0, 100 mM NaCl, 1 mM EDTA, 0.5 mM EGTA, 0.1% Na-Deoxycholate, 0.5% N-lauroylsarcosine, 1X protease inhibitors) on Biorupter^®^ (Diagenode) by two rounds of 10min sonication to obtain 300-600bp range chromatin fragments. The resulting sheared chromatin was immunoprecipitated (IP) using 20 μg of antibody against POL2 (Santa Cruz, N-10), a non-specific Isotype Mouse IgG as a mock control (Santa Cruz). The IP complexes were then extracted using Protein A/G Dynabeads and washed five times using RIPA washing buffer (50 mM HEPES-KOH, pKa 7.55, 500 mM LiCl, 1 mM EDTA, 1.0% NP-40, 0.7% Na-deoxycholate). The DNA was extracted from the beads using an elution buffer (50 mM Tris-HCl, pH 8.0, 10 mM EDTA, 1.0% SDS) and quantified by qPCR using primers designed to amplify the promoter region of the reporter construct. Information on primer sequences is described in Supplementary Information 1.

### Fluorescent microscopy and live imaging

1 μg of pmCGFP reporter constructs containing 0, 1, 2 copies of TCTCGCGAGA along with a CMV promoter-driven Blue Fluorescent Protein expression plasmid (CMV-BFP) were transfected into HEK293T cells. Images were taken 24 h post-transfection using a Nikon Eclipse TE2000-E fluorescence microscope. For live imaging, images were taken every 15 min with an exposure time of 1 sec immediately after reporter transfection for 24 h in an incubating chamber supplied with humidity and 5 percent conditions. 16-bit Tiff images from individual channels were used to generate MOV files using the Videomach software (http://gromada.com/videomach/). The final production video was produced using Adobe Premiere CC 2017.

### Double nickase Cas9 mediated genome editing of DENR promoter

Short guide RNAs (sgRNAs) to target *DENR* promoter were designed using the MIT CRISPR sgRNA design tool (<u>http://crispr.mit.edu/</u>). The DNA sequences of guides (Supplementary Information 1) were then cloned into pSpCas9 (BB)-2A-GFP (PX458) and pSpCas9 (BB)-2A-Puro (PX459) V2.0 (Addgene plasmid numbers 48138 and 62988). Constructs were then co-transfected into HEK 293T cells and 24 hrs later selected for Puromycine resistance (3 μg/mL) for another 72 hours. GFP-expressing single cells were sorted using Aria II FACS instrument and incubated in 96 well dishes for two weeks to form visible cellular clones. DNA was extracted from the clones using QuickExtract™ solution (Epibio), and successful deletions were confirmed by Sanger sequencing of PCR products. Ribbon sequences were produced using the pyRibbon software which we deposited in https://github.com/AminMahpour/pyRibbon.

### Immunoblotting

Cells were lysed in NET-N buffer (100 mM NaCl, 20 mM Tris-HCl pH 8.0, 0.5 mM EDTA, 0.5% NP-40) supplemented with protease inhibitors cocktail at 4°C. In all experiments, 20 μg of total proteins/lane analyzed by SDS-PAGE followed by blotting as described in Previs, et al. ^52^. Antibodies included those specific for DENR (Santa Cruz 22), GFP (Santa Cruz B-2), mCherry (Abcam 1C51) or alpha-tubulin (Santa Cruz A-6) as a loading control.

### Oligonucleotide pull-down assay

To determine whether CGCG elements can bind to Kaiso, we separately synthesized biotin-tagged DNA duplex that contained unmodified TCTCGCGAGA, TCTC<u>T</u>CGAGA or completely methylated (TCT^me^CG^me^CGAGA). 10 μM from each duplex were bound and washed to 100 μl Streptavidin Dynabeads as recommended by the manufacturer (Invitrogen). HEK293T cells were lysed using NET-N buffer containing protease inhibitors cocktail (Sigma) and incubated on ice for 30 min. Lysates were centrifuged at 12000 RPM for 10 min to pellet cellular debris, and supernatant representing 500 μg protein was mixed with duplex-charged beads and incubated at 4°C overnight. The beads were washed five times with NET-N buffer, incubated with 50 μl Laemmli loading buffer (1X: 0.02% w/v bromophenol blue, 4% SDS, 20% glycerol, 120 mM Tris-Cl, pH 6.8) and boiled for 5 min to elute bound proteins. The proteins were analyzed by immunoblotting for Kaiso (Santa Cruz D-10) and control antibody.

### Rapid Amplification of cDNA ends (5′ RACE)

To determine divergent TSSs, we transfected near confluent HEK293T cells in 10 cm dishes with 5 μg LuBiDi construct along with 0.5 μg pEGFP-C1 to monitor transfection in the following day. RNA was extracted as described before 72 h after transfection. The quality and purity of RNA were evaluated using Agilent 2100 Bioanalyzer and samples with RNA integrity number (RIN) values >= 8.0 were selected for further analysis. The SMARTer 5´ RACE (Clontech) protocol was used to determine divergent TSSs from 10 μg of total RNA. Briefly, the RNA was first reverse transcribed at 42°C for 90 min using poly-dT primers and extended beyond TSS using RT-mediated template switching that employs the SMARTer IIA Oligonucleotide as the template only when the 5′ cap is encountered. The resulting cDNA products were amplified using specific internal primers for either firefly or Renilla plus the Clontech Universal Primer Mix (UPM). A GFP primer set was used as an internal control. Primer sequences used in RACE experiments are provided in the Supplementary Information 1. The PCR products containing TSS were directionally cloned into the linearized pRACE vector using the Infusion HD system, and individual bacterial clones were obtained following transformation of the ligated products into Stellar competent cells. Sanger sequencing of the resulting plasmid clones (using M13 primer) was used to identify TSSs.

### Genomic analysis

#### Motif Discovery

The CpG island annotation track in the human genome (hg38) was downloaded from the UCSC genome browser (https://genome.ucsc.edu), and sequences that overlap with K562 DNase-seq peaks track were extracted using Bedtools ^53^. The resulting sequences were used for motif discovery using the findMotifgenomewide script in the Homer bioinformatics software suite using default command line arguments for the human genome ^54^.

#### Genomic annotation and Metagene analysis

The scanMotifgenomewide script from the Homer program version 4.8 was used to locate all instances of motif 7 and 10 in human (hg38) and mouse (mm9) genomes. The annotatePeaks script (Homer) was used to identify motif co-occurrence, genomic annotations, metagene, and enrichment analysis.

#### ENCODE Conservation, DNase-seq, GRO-Cap, WGBS

Processed data points for hg38 were extracted and processed using Wigman software for 50 bp upstream and downstream windows for each motif occurrence. For ENCODE WGBS (accession number ENCFF867JRG). The PhyloP and PhastCons conservation scores for hg38 assembly were downloaded from the UCSC genome browser (http://hgdownload.cse.ucsc.edu/downloads.html). ENCODE accession number ENCFF867JRG was used for K562 DNase-seq data. The GRO-Cap dataset for K562 and GM12878 cell lines with GEO accession number of GSM1480321 was used to analyze nascent transcripts in promoters. POL2 ChIP-seq from K562 cell line with the accession number of ENCFF000YWS was used to determine POL2 occupancy state on CGCG elements. Heatmap plots were generated using the in-house written Wigman software (https://github.com/AminMahpour/Wigman).

#### Gene Ontology and gene network analysis

Bedtools Closest feature was used to compile a list of genes that their annotated TSS are less than 500bp from CGCG elements on both plus and minus strand from the latest hg38 GTF annotation file (http://www.ensembl.org/info/data/ftp/index.html). A custom script was written and used to determine the number of CGCG elements in annotated coding, non-coding, uni-and bi-directional CGI promoters.

Gene Ontology (GO) analysis performed using GOrilla gene enrichment analysis platform. A list of CpG islands-associated genes was used as the background genes for enrichment analysis ^55^. GO enrichment score is defined as (***b***/***n***)/(***B***/***N***) where N is the total number of background CpG island-associated genes that have a GO term, B is the number of genes associated with a specified GO term, and n is the number of genes whose promoter contain CGCG element and b is the number of genes in the intersection. Gene set interaction networks were generated and analyzed using REACTOME package v53 (<u>http://www.reactome.org/</u>). Network were visualized graphically using Cytoscape software version 3.5 (<u>http://www.cytoscape.org/</u>).

#### Start-seq analysis

Start-seq from mouse bone-marrow derived macrophages was published previously and is available for download from GEO website (GSE62151, https://www.ncbi.nlm.nih.gov/geo/). Data were analyzed as described previously. Briefly, reads were aligned uniquely to the mm9 genome allowing a maximum of two mismatches with Bowtie version 0.12.8 (-m1 -v2). Sense and divergent TSS were assigned as defined previously. Start-seq heat maps depict Start-RNA reads in 10 bp bins at the indicated distances with respect to the TSS. Heatmap plots were generated using Partek Genomics Suite version 6.12.1012.

Individual CGCG element occurrences were identified with FIMO ^56^. A ±1 kbp window around TSSs was scanned with a position weight matrix for the CGCG motif with a p-value threshold of 0.001. Motif occurrences were mapped with respect to TSS locations using custom scripts and counted in 10-mer bins. Composite Metagene distributions were generated by summing motifs at each indicated position with respect to the TSS and dividing by the number of TSSs included within each group.

### Statistical Analysis

All plots were generated and analyzed using GraphPad Prism version 7. Unless noted otherwise, all statistical analyses were performed using Student t-test. The following p-values are presented as *: p<0.05, **: p<0.01, ***: p<0.001, ****: p<0.0001. Error bars represent standard deviation (S.D) from the mean.

## Author Contributions

A.M conceived the project, performed experiments, analyzed experimental and informatics data, interpreted the results and wrote the manuscript. B.S analyzed mouse Start-seq data and assisted in manuscript preparation. D.S and I.G contributed to the project design and edited the manuscript. T.O secured funding and supervised the project.

## Acknowledgment

Authors would like to thank William Burhans for his valuable input and criticism of the manuscript, and for members of the Ouchi and Gelman laboratories for discussion. This work was supported in part by NIH CA90631 (T.O.) and Susan Komen Breast Cancer Foundation (T.O.). The first author would like to dedicate this paper to his parents.

## Supplementary Figures

**Figure 1.**
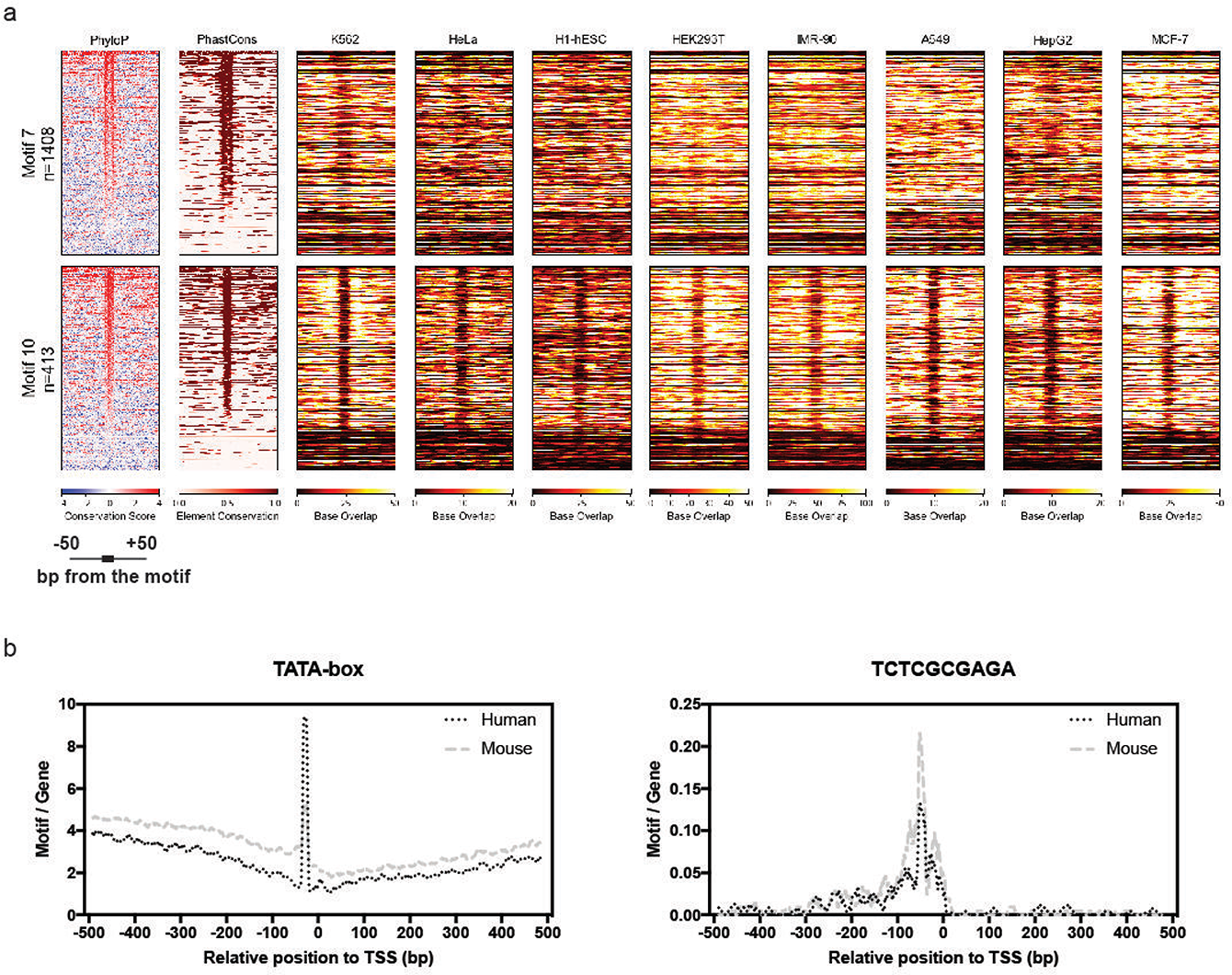
The CGCG element (motif 10) is associated with DNase-seq footprint in different cell lines. a) ENCODE DNase-seq footprints of motif 7 and 10 for available cell lines. b) TCTCGCGAGA motif occurs within 50bp of annotated TSSs in the human and mouse genomes.

**Figure 2.**
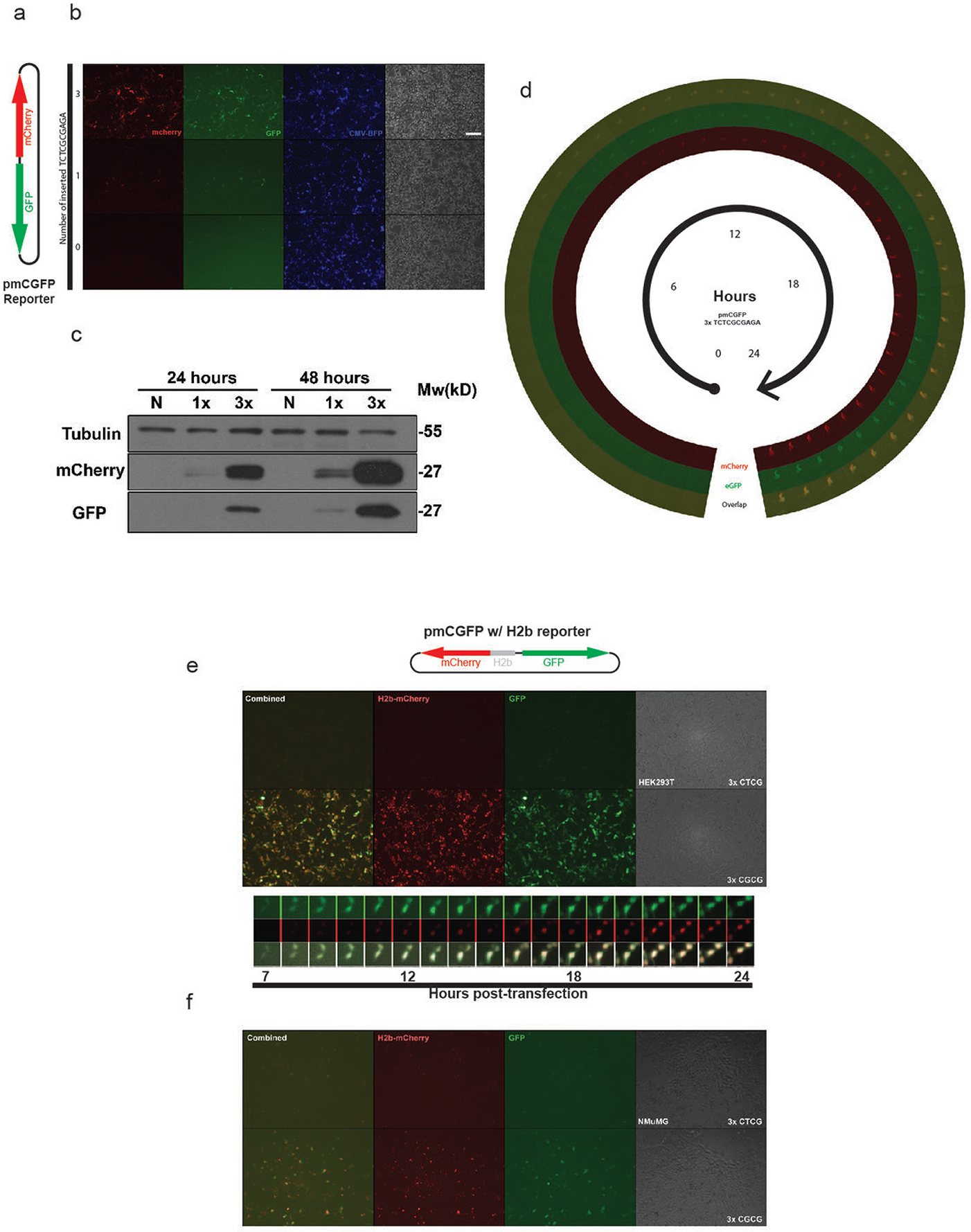
The CGCG element promote simultaneous expression of GFP and mCherry genes in the pmCGFP reporter construct. a) The pmCGFP bidirectional reporter structure. b) The fluorescence image of HEK293T cells transfected with pmCGFP constructs containing 0, 1 or 3 copies of TCTCGCGAGA motif after 24 hours. CMV-driven BFP expression was used as an internal control c) Immunoblots showing levels of GFP and mCherry expression 24 and 48 hours post transfection. d) Time-lapse imaging of HEK293T cells transfected with the pmCGFP containing three copies of TCTCGCGAGA for 24 hours shows that both reporters are simultaneously expressed few hours after transfection. Scale bar is 100 μm. e) CGCG element confer bidirectional expression of GFP and H2b-mCherry reporter genes in HEK293T. Time-Lapse images of HEK293T cell line transfected with a pmCGFP-H2b (h2b-mcherry fused gene) reporter. Please note delayed H2b-mCherry signals as the fused mCherry protein is being trafficked into the nucleus. f) Images of NMuMG mouse cells transfected with pmCGFP-H2b construct containing either 3 copies of wild-type TCTCGCGAGA motif or 3 copies of TCTC<u>T</u>CGAGA mutant motif.

**Figure 3.**
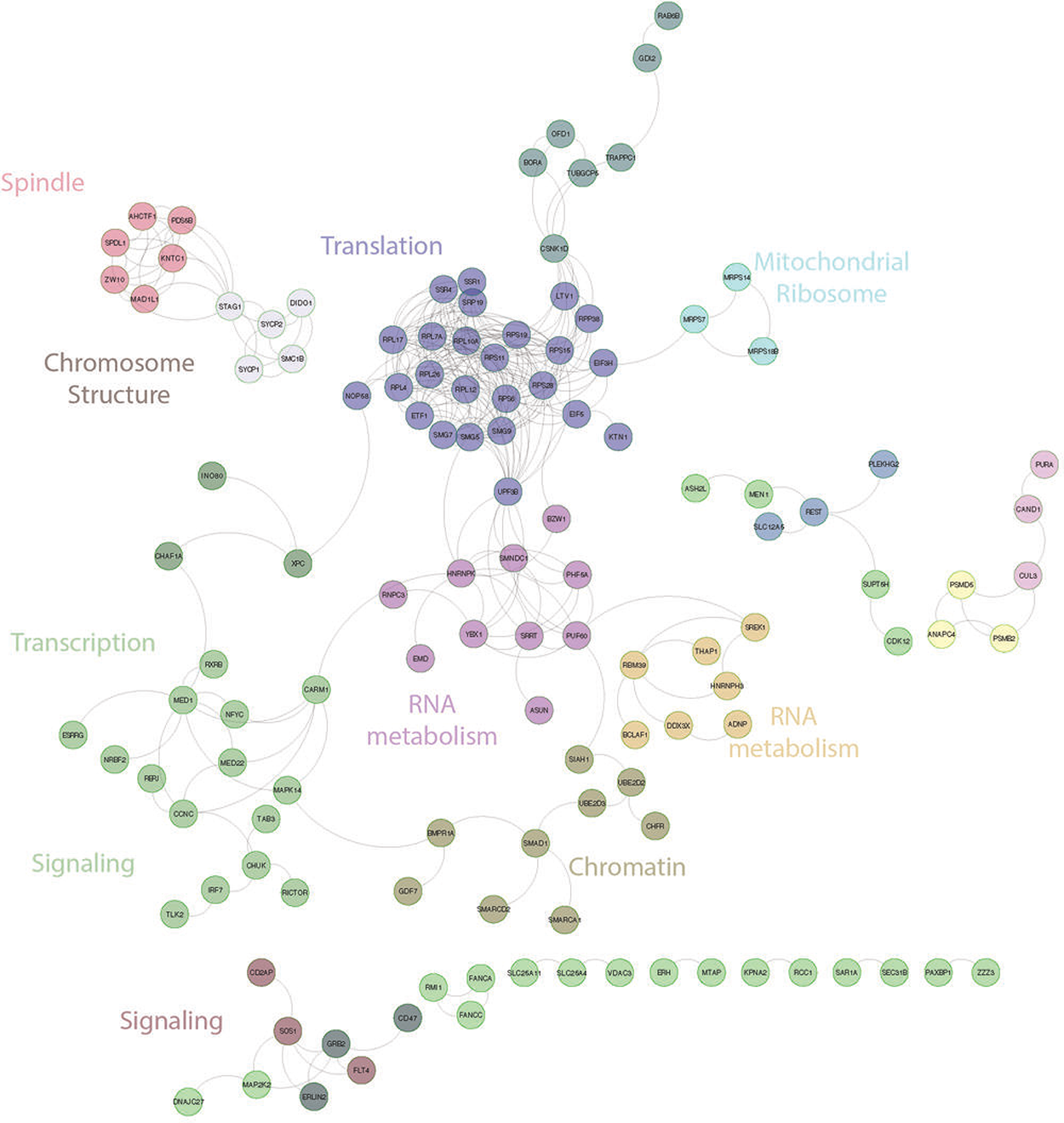
REACTOME Interaction network analysis of CGCG containing promoters. An analysis of genes that contain CGCG elements in their promoters found that most of these genes can be clustered into distinct functional groups as indicated in the figure.

**Figure 4.**
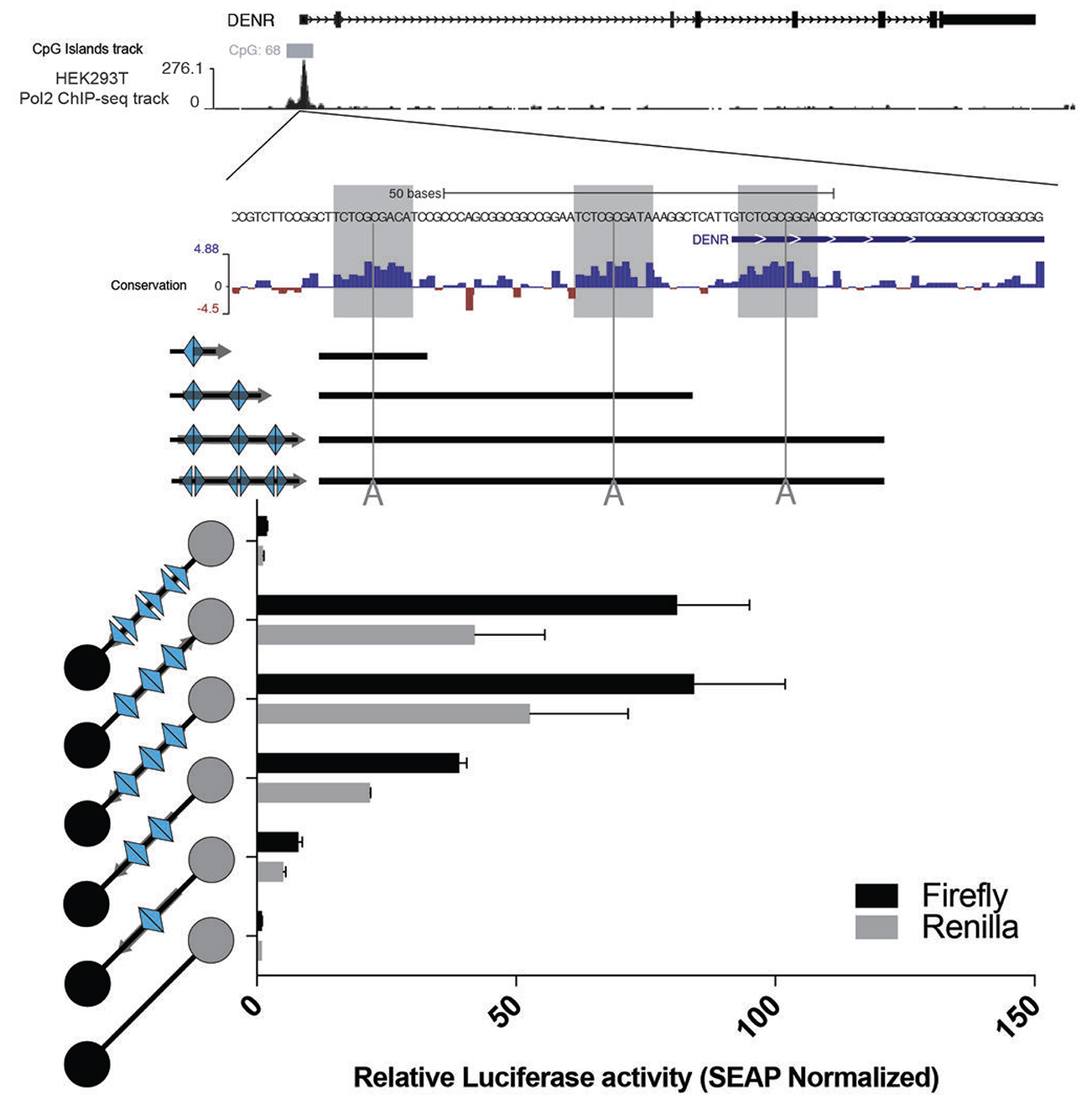
CGCG elements in the DENR promoter promote divergent transcription. The CGCG elements in the *DENR* promoter, regardless of the insert direction, activated bidirectional reporter genes. Insertion of an “A” in the center of CGCG elements eliminated the promoter activity.

**Figure 5.**
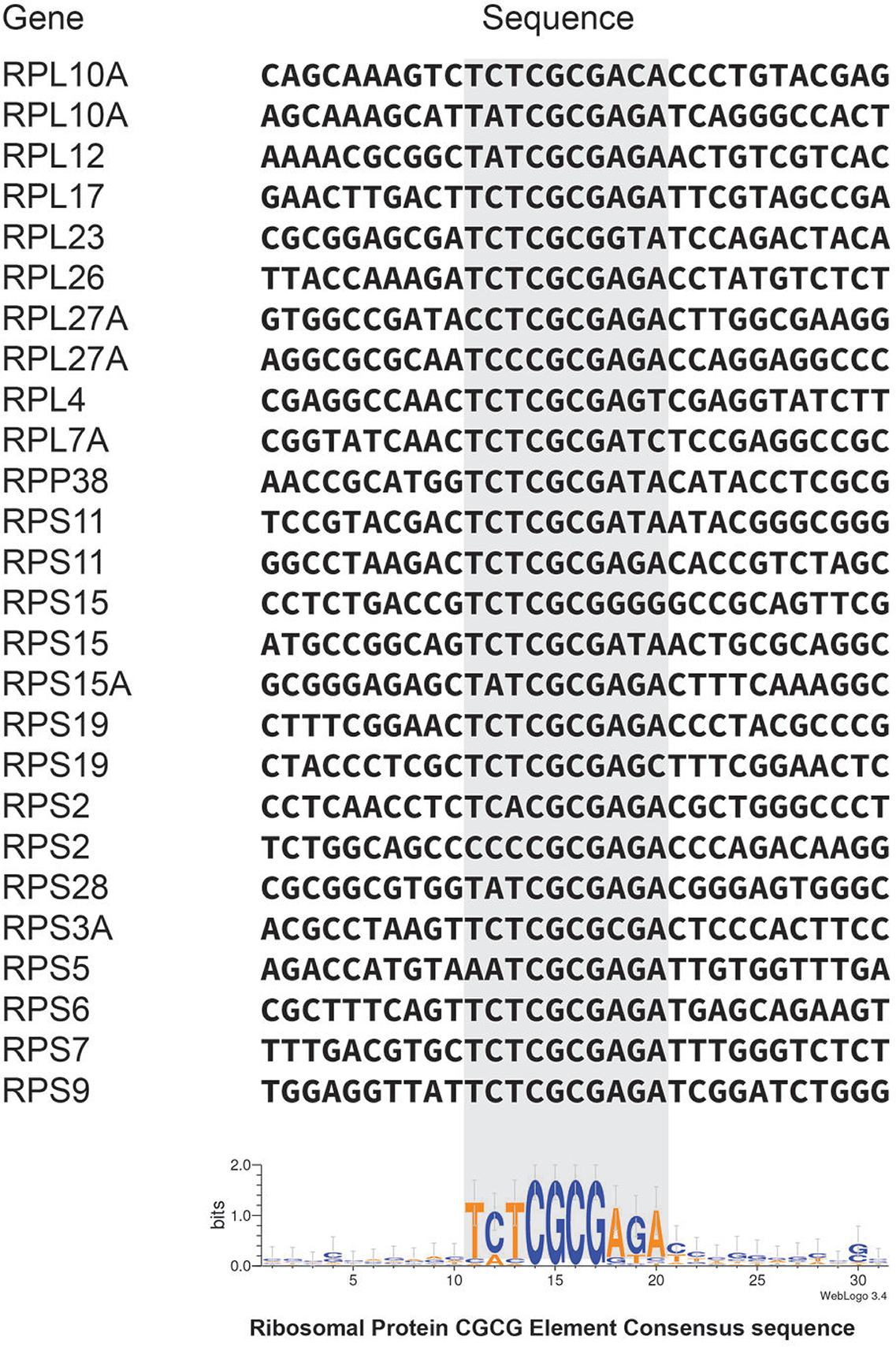
CGCG elements are enriched in Ribosomal protein promoters. Aligned sequences of CGCG elements and flanking regions in the promoters of Ribosomal proteins genes. Ribosomal genes that contain CGCG element are devoid of TCT motif.

## Supplementary video

TCTCGCGAGA motif activated the simultaneous expression of both GFP and mCherry fluorescent reporters.

